# On-scalp MEG system utilizing an actively shielded array of optically-pumped magnetometers

**DOI:** 10.1101/471391

**Authors:** Joonas Iivanainen, Rasmus Zetter, Mikael Grön, Karoliina Hakkarainen, Lauri Parkkonen

## Abstract

The spatial resolution of magnetoencephalography (MEG) can be increased from that of conventional SQUID-based systems by employing on-scalp sensor arrays of e.g. optically-pumped magnetometers (OPMs). However, OPMs reach sufficient sensitivity for neuromagnetic measurements only when operated in a very low absolute magnetic field of few nanoteslas or less, usually not reached in a typical magnetically shielded room constructed for SQUID-based MEG. Moreover, field drifts affect the calibration of OPMs. Static and dynamic control of the ambient field is thus necessary for good-quality neuromagnetic measurements with OPMs. Here, we describe an on-scalp MEG system that utilizes OPMs and external compensation coils that provide static and dynamic shielding against ambient fields.

In a conventional two-layer magnetically shielded room, our coil system reduced the maximum remanent DC-field component within an 8-channel OPM array from 70 to less than 1 nT, enabling the sensors to operate in the sensitive spin exchange relaxation-free regime. When compensating field drifts below 4 Hz, a low-frequency shielding factor of 22 dB was achieved, which reduced the peak-to-peak drift from 1.3 to 0.4 nT and thereby the standard deviation of the sensor calibration from 1.6% to 0.4%. Without band-limiting the field that is compensated, a low-frequency shielding factor of 43 dB was achieved.

We validated the system by measuring brain responses to electric stimulation of the median nerve. With dynamic shielding and digital interference suppression methods, single-trial somatosensory evoked responses could be detected. Our results advance the deployment of OPM-based on-scalp MEG in lighter magnetic shields.

## 1. Introduction

Magnetoencephalography (MEG) is a non-invasive neuroimaging technique that measures the magnetic fields of electrically active neuron populations in the human brain (Hämäläinen et al., 1993). Spatial sampling of the neuromagnetic field with an array of hundreds of sensors allows localisation of brain activity with millimeter accuracy in favorable conditions. The frequency content of the neuromagnetic field mostly lies in a band from 1 to 80 Hz while its amplitude ranges from femto-to picotesla (Baillet, 2017). The detection of the weak neuromagnetic field thus necessitates highly sensitive magnetometers. In addition, the measurement must be shielded from external magnetic disturbances, such as Earth’s magnetic field and its fluctuations, and fields generated by power lines and moving magnetic objects, as the interfering fields often lie in the frequency band of interest, and their amplitudes can exceed those of the neuromagnetic fields by several orders of magnitude (Taulu et al., 2014).

Until recently, MEG systems have mostly employed superconducting quantum interference devices (SQUIDs); a typical SQUID sensor of a commercial MEG device has a sensitivity of about 3 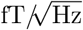 and a pick-up loop surface area of about 4 cm^2^. Sufficient magnetic shielding for the measurement has been usually achieved by performing the measurements inside a large magnetically shielded room (MSR) comprising layers of highly permeable and conductive materials.

Liquid helium cooling of the SQUIDs to their critical temperature requires thermal insulation between the SQUID and the subject’s head, limiting the measurement distance of the neuromagnetic fields to about 2 cm from the scalp. Novel magnetometers, such as optically-pumped magnetometers (OPMs) (Budker and Romalis, 2007) and high-T_c_ SQUIDs (Faley et al., 2017), have lately emerged to rival SQUIDs in biomagnetic measurements. In contrast to SQUIDs, these new sensor types allow measurement of the neuromagnetic field within millimeters of the subject’s scalp, increasing the measured signal amplitudes as well as the spatial resolution and the information content of the measurement (Boto et al., 2016; Iivanainen et al., 2017; Riaz et al., 2017).

OPMs are based on detecting the Larmor precession of spin polarization that is generated into an alkali-atom vapor by means of optical pumping. The sensitivity of an OPM can be brought to a sufficiently high level for MEG by operating it in the spin-exchange relaxation-free (SERF) regime that is achieved when the alkali-atom vapor density is high and the absolute magnetic field is very low (Allred et al., 2002). To date, numerous SERF OPMs with a sensitivity and size suitable for whole-scalp MEG have been presented (Colombo et al., 2016; Sheng et al., 2017a; Osborne et al., 2018; Sheng et al., 2017b). In addition, multichannel OPM-based systems have also been demonstrated. Alem and colleagues (2017) have presented an OPM system comprising 25 microfabricated sensors and demonstrated its use with a phantom. Borna and colleagues (2017) showed a 20-channel OPM-MEG system operating inside a person-sized cylindrical shield, while Boto and colleagues (2018) demonstrated a wearable OPM-based MEG sensor array acting inside a standard MSR for SQUID-MEG.

The magnetic shielding requirements for OPMs especially at near-DC frequencies present challenges when this new technology is adopted to practical MEG use. The SERF condition implies that the magnetic field has to be very low (practically below 1 nT; ideally ~0 nT). OPMs usually have small sensor-wise coils that null the DC field within the sensitive volume of each sensor. For example, such a coil set in one commercial OPM can compensate fields up to 50 nT within its volume (Osborne et al., 2018), however, the remanent DC field inside an MSR may exceed this value (see e.g. Johnson et al. (2013)). In addition, nT-level field variations inside the volume of the sensor affect the sensor calibration by altering the gain and tilting the sensitive axis of the sensor (see Appendix A). Such field variations can result from at least two distinct mechanisms. First, external interference may leak in to the MSR and generate a temporally and spatially varying residual field in addition to the DC remanent field within the room. Second, sensor movement (translation and rotation) in that field can also change the field experienced by the sensor. These field changes are especially prominent when the sensor array is wearable: the sensors move together with the subject’s head inside the shield. Carefully nulling the static remanent field inside the volume of the movement reduces such field changes. For example, for their wearable OPM array, Boto and colleagues (2018) nulled the static remanent field with three coils producing homogeneous fields and two gradient coils by using the output of four reference sensors that were kept away from the sensors that measured brain activity.

To address the aforementioned issues due to static and dynamic components of the residual field inside a shielded room, we have designed and constructed an active magnetic shielding system that is used with a multichannel OPM-based MEG sensor array. The shielding system comprises eight coils and feedback electronics, achieving two goals. First, it can null the static remanent field (homogeneous components up to ~160 nT; gradients up to ~100 nT/m) to bring the OPMs to their dynamic range together with the sensor-wise smaller coils. Second, it actively nulls the field within the sensor array using a negative feedback loop and thus reduces OPM calibration drifts due to the temporal field drifts inside the MSR. The static compensation and the feedback loop can be driven with a variable number of sensors and with different sensor configurations so that they can be adjusted to the given measurement and sensor set-up. In addition, the feedback loop incorporates an adjustable low-pass filter with a very low cut-off frequency so that the same sensors can be used for compensating low-frequency field drifts as well as for registering higher-frequency brain responses. Here, we describe our OPM-based on-scalp MEG system and the shielding performance of its coil system in a two-layer MSR. We also quantify OPM calibration drifts in the MSR with and without dynamic shielding. Last, we present an on-scalp MEG measurement of neuromagnetic responses to electric stimulation of the median nerve by employing our system with dynamic shielding.

## 2. Materials and methods

### 2.1. Characteristics of the shielded room

In this section, we shortly present the characteristics of the two-layer shielded room in Aalto University (MSD-2S, ETS-Lindgren Oy, Eura, Finland) and motivate the construction of the shielding system.

The remanent field inside the two-layer MSR near the center of it is presented in Fig. 1. The field was measured with a three-axis fluxgate magnetometer (Mag-03MC1000, Bartington Instruments, Oxford, UK) in a volume of (19.2 cm)^3^. The field amplitude was 59–84 nT while the homogeneous field components were −10, 40 and −60 nT along the shorter and longer walls of the room and the vertical axis (*x, y* and *z*, respectively). The maximum field gradient was about 60 nT/m. Given the upper limit of 50 nT for the internal DC field compensation of the aforementioned commercial OPM, the remanent field must be nulled with coils external to the sensor.

**Figure 1:**
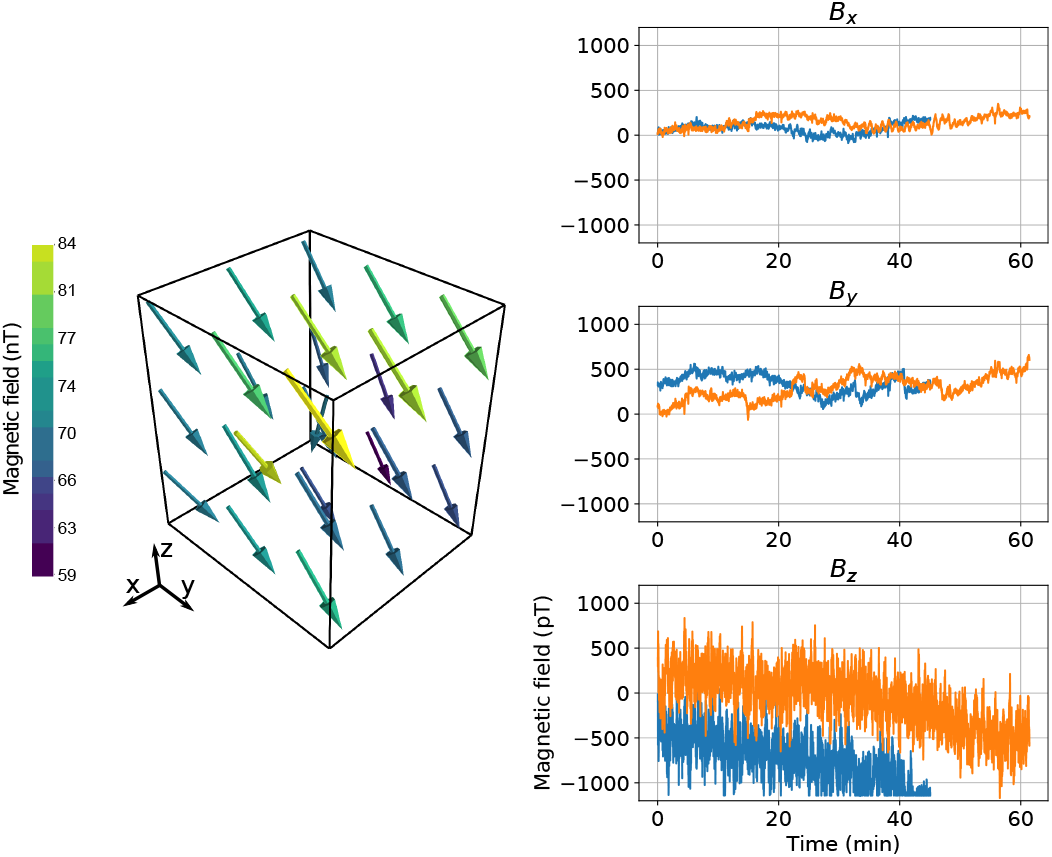
Magnetic field inside the two-layer magnetically shielded room of Aalto University. **Left:** Quiver plot of the remanent DC field measured with a three-axis fluxgate magnetometer in a volume of (19.2-cm)^3^ near the center of the shielded room. The measurement points are distributed uniformly in a 3 × 3 × 3 grid in the measurement volume. Color indicates the amplitude of the magnetic field (scale in nT). The shorter and longer walls of the room are along x and y axes while z denotes the vertical axis. **Right:** Drifts of the three orthogonal components of the magnetic field from two independent measurements (blue/orange) measured with OPMs. In the other measurement (blue), *B_z_* was clipped due to exceeding the range of the analog-to-digital converter.

Even if the internal coils of the sensor could null the field locally, global nulling is beneficial as small sensor movements in a large remanent field would produce large measurement artefacts and calibration errors. For example, even at the center of our shielded room, rotation of the sensor by 1° causes field variation up to 0.7 nT and moving the sensor by 0.5 cm produces an artefact with an amplitude of 0.3 nT, which is three orders of magnitude larger than brain responses.

In addition to the measured DC fields, Fig. 1 shows temporal drifts of the magnetic field components inside the MSR as measured with an OPM (QuSpin Inc., Louisville, CO, USA). The field drifts in the *z*-direction much more rapidly than in the *x*- and *y*-directions: during ten seconds, the field can drift 130 and 190 pT in the x- and y-directions and as much as 1 nT in the *z*-direction. The peak-to-peak field excursions during about an hour are approximately 370, 720 and 2000 pT for the *x*-, *y*- and *z*-components of the field, respectively.

During a two-year observation period, the remanent DC field inside the room has been changing; the maximum DC field component at the center of the room has been about 200 nT. We suspect that such variation is due to the magnetization of the walls of the MSR caused by prepolarization pulses applied in the MEG–MRI system (Vesanen et al., 2013) that is sharing the shielded room.

The compensation system should be thus designed so that it is capable of zeroing the DC remanent field (homogeneous components and gradients) and its temporal variations inside this room. The long-term variation in the remanent field should be taken into account when deciding the gains of the external field coils.

### 2.2. Overview of the system

Figure 2 provides an overview of our system. The system consists of eight rectangular coils that are mounted on the sides of a (78-cm)^3^ cube, OPMs positioned in a helmet located near the center of the coil cube and electronics for data acquisition, OPM control as well as for calculating and feeding feedback currents to the coils.

**Figure 2:**
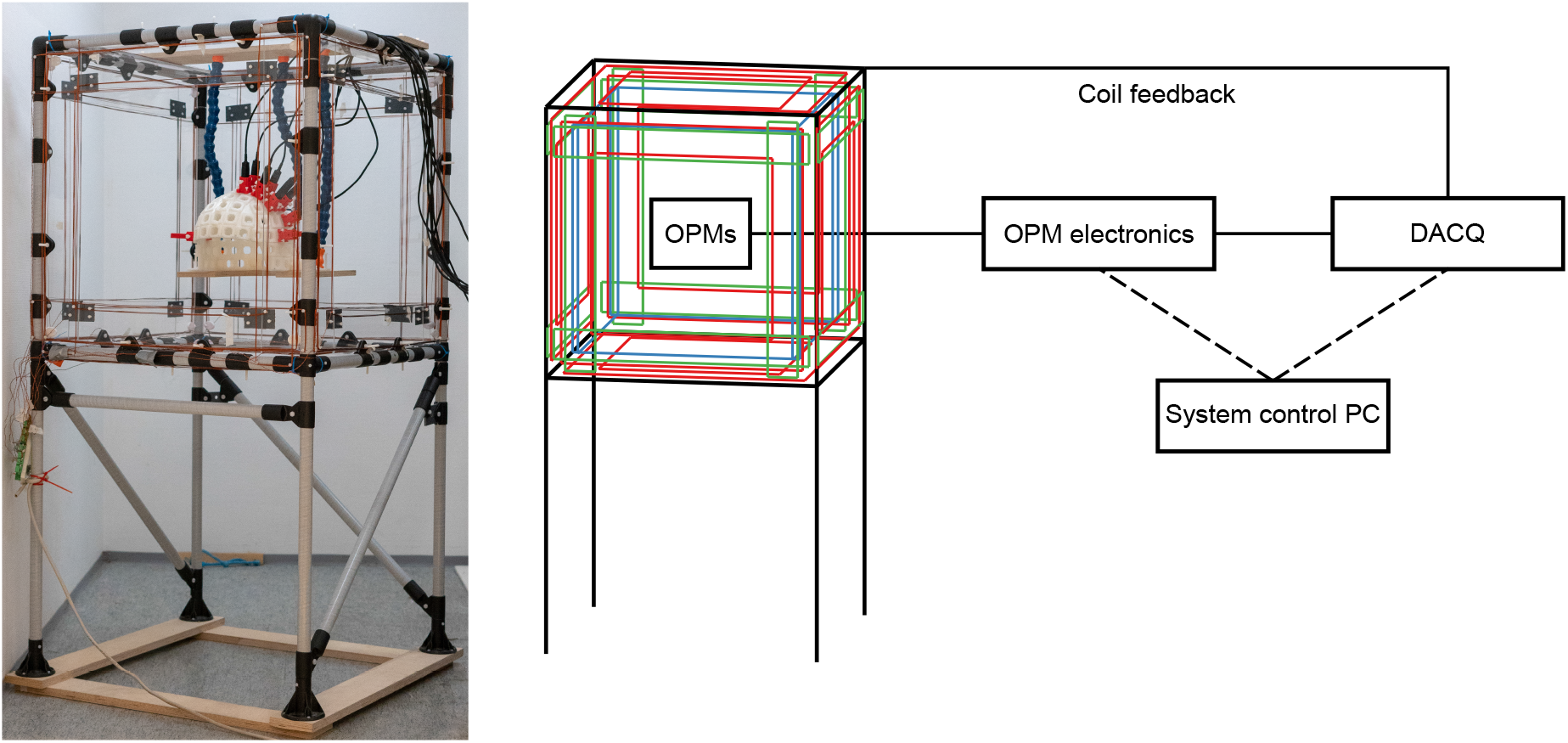
Overview of the active shielding system. **Left:** Picture taken from the system in a shielded room with OPMs attached to the sensor helmet. **Right:** A schematic figure illustrating the main components of the system. One PC is used to control the OPM electronics and the electronics for data acquisition and computing feedback currents (DACQ).

Our system uses OPMs produced by QuSpin Inc. (Gen-1.0 QZFM; Osborne et al. 2018). The OPMs are placed in a helmet that is located in the center of the coils and that is attached to the shielding system with three flexible support hoses (Loc-Line, Lake Oswegon, OR, USA); see Fig. 2. The helmet has 102 locations for the sensors while the flexible support allows adjustment of the position of the helmet inside the system. The measurements in this paper are performed with eight OPMs although the system is scalable to more sensors.

The OPM sensor head comprises a resistively-heated vapor cell containing ^87^Rb atoms and a mixture of buffer gases, single-mode vertical cavity surface emitting laser (VCSEL), photodiode for measuring transmitted light intensity, optical components for controlling the light beam and its path, and coils for nulling the DC field and for providing field modulation. The rubidium atoms are spin-polarized using circularly-polarized light from the VCSEL. The generated spin polarization undergoes Larmor precession in the applied magnetic field, which in turn modifies the absorption of the light that polarized the atoms; the light absorption detected with the photodiode then serves as an indicator of the magnetic field amplitude and has a Lorentzian lineshape around zero field. To get the sensor into the SERF regime, the vapor cell is heated to about 150 ° C and the field coils are used to null the field inside the vapor cell. In addition, to make the sensor sensitive along a specific axis and to suppress technical noise, the magnetic field inside the vapor cell is modulated along that axis at 923 Hz with the field coils. The photodiode signal is then demodulated with a lock-in amplifier to yield the output signal of the sensor that has a dispersive lineshape around zero field.

The OPM sensor is housed in a 13 × 19 × 110 mm^3^ package. The sensitivity of the sensor is around 7–13 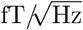 while the sensor bandwidth is about 135 Hz. By selecting the internal coils providing field modulation, the sensitive axis of the OPM can be chosen along the longitudinal or transverse direction with respect to the sensor housing in the single-axis mode or along both in the dual-axis mode; however, the sensitivity decreases in the dual-axis mode. The internal coils of the sensor can zero fields up to 50 nT. For a more comprehensive review of the sensor design and performance, see Osborne et al. (2018).

Each OPM sensor has its own control electronics (QuSpin Inc.) that drives the laser, controls the temperature of the vapor cell, digitizes the photodiode output, performs the lock-in detection, and feeds current to the internal DC-field-zeroing coils (Osborne et al., 2018). The OPM electronics are controlled digitally with a custom software on the measurement PC that is written in the Python programming language. The electronics provide the OPM output to the PC in both analog and digital format.

We collect the analog output of the OPM electronics with a data-acquisition system (DACQ) that is based on the electronics of a commercial MEG system (MEGIN Oy (formerly Elekta Oy), Helsinki, Finland). The DACQ consists of digital signal processor (DSP) units that digitize and collect the analog OPM output, compute coil feedback current and feed the coils. In addition to the feedback currents, the DACQ can also output DC, sinusoidal, triangular and white-noise currents to the coils. The DACQ can be controlled in real time over Ethernet using a proprietary protocol on a TCP/IP (transmission control protocol / internet protocol) connection with an interface in Python. The electronics and software are described in more detail in Sec. 2.2.2.

#### 2.2.1. Compensation coils

We designed the coil system to achieve full control of the homogeneous components and first-order gradients of the magnetic field. One of Maxwell’s equations 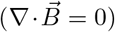 restricts the number of independent longitudinal gradients to two while another Maxwell’s equation 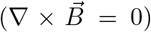 limits the number of independent transverse gradients to three. Thus, for the complete control of the homogeneous fields and first-order gradients, three homogeneous-field, two longitudinal-gradient and three transverse-gradient coils are necessary. We designed coils that produce three homogeneous fields along the three orthogonal axes of the cartesian coordinate system 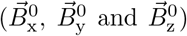, longitudinal gradients along *x* and *y* (*∂B*_x_/*∂x* and *∂B*_y_/*∂y*) and transverse gradients along *y* and *z* (*∂B*_x_/*∂y, ∂B*_x_/*∂z* and *∂B*_y_/*∂z*).

The coils were designed to fit on the sides of an approximately (78-cm)^3^ cubical volume. The dimensions were determined by the width of the door frame and the space available in the two-layer MSR that also houses the MEG–MRI system. Further, the coils were designed such that they allow relatively easy access for the subject and do not severely limit the subject’s visual field. We selected rectangular coil designs due to their ease of construction. The coils were designed with a custom MATLAB (Mathworks Inc., Natick, MA, USA) program that calculates the magnetic field of an arbitrary arrangement of straight wire segments. For the homogeneous-field coils, the figure of merit for the design was the homogeneity of the magnetic field component over a volume of (20 cm)^3^ at the center while that for the gradient coils was the gradient-field linearity in that same volume. Field homogeneity was defined as the relative deviation of the magnetic field component in the volume to that at the center while the gradient-field linearity was defined as the relative deviation from a linear field profile.

Fig. 3 shows the coil designs together with their computed fields in a centrally-aligned (20-cm)^3^ volume. For the homogeneous-field coils, we adopted the design by Ditterich and Eggert (2001) where the authors had a similar target; they had to optimize the magnetic field homogeneity of a rectangular coil when space was limited and access into the middle of the coils was required. The homogeneous-field coil consists of two sets of coaxial square coils with sidelengths of 70.0 and 52.5 cm, separated by 70.0 cm. The currents in the larger and smaller coils flow in opposite directions and the ratio of the current amplitude in the smaller coil to that in the larger coil should be 0.58. We implemented this by connecting both the larger and smaller coil in series and winding them in opposite directions so that the number of turns for the larger coil was 14 and for the smaller coil 8 (ratio: 0.57). Our calculations indicated that this design should yield a magnetic field homogeneity of 2.0% within the (20-cm)^3^ volume. The number of turns in the *B*_x_ and *B*_y_ homogeneous field coils are the same (14 and 8) while the *B*_z_ coil has twice as many turns (see Sec. 4.2).

**Figure 3:**
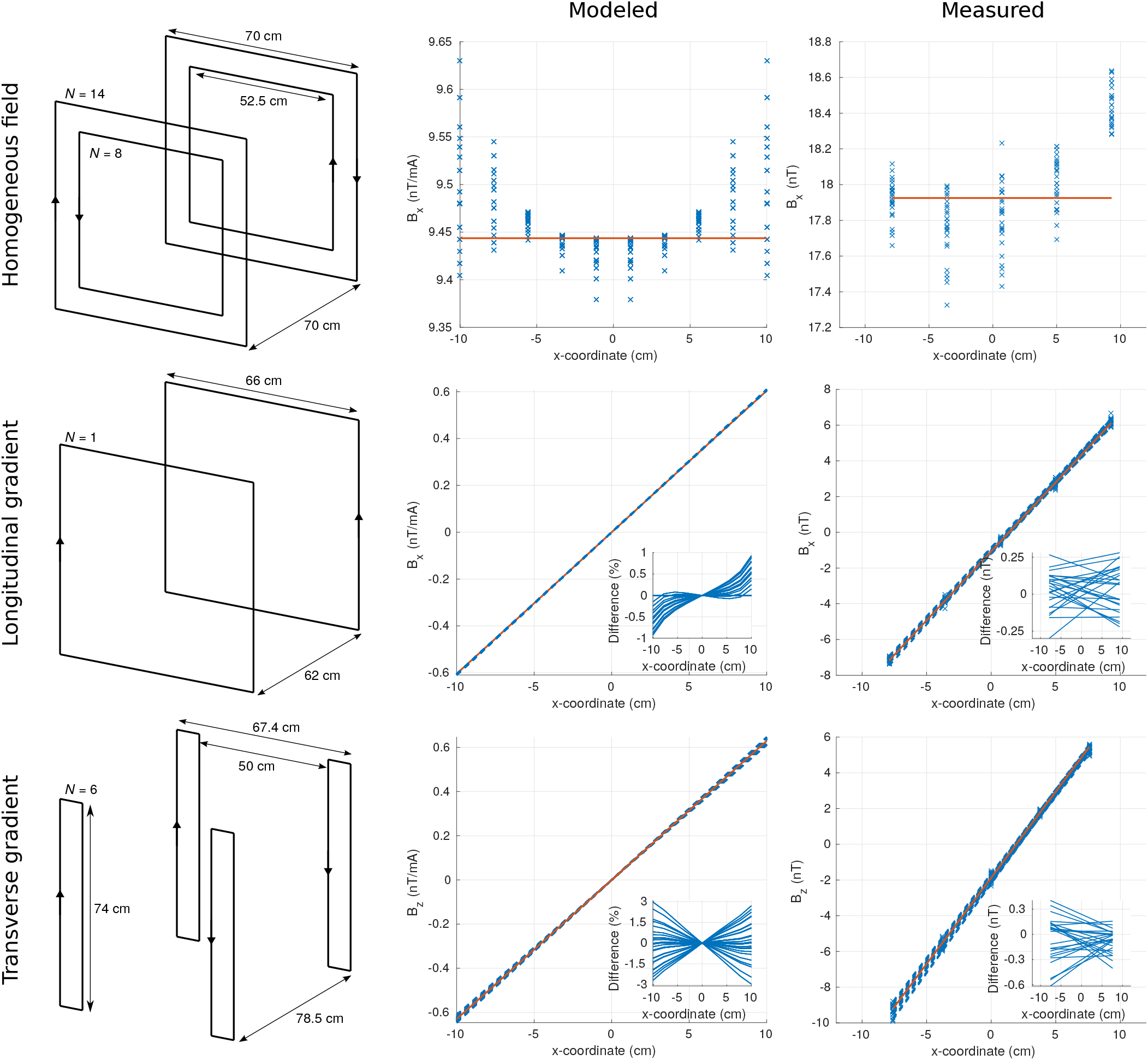
The designs (**left**) of the coils in the active shielding system together with their modeled (**center**) and measured (**right**) fields. The measured field values are marked with crosses while the dashed lines present the estimated slopes of the field components. The solid lines indicate the desired homogeneous components or gradients. The insets show the difference to the desired slopes.

Square Maxwell coil design was used for the longitudinal gradient coils: two coaxial square coils with side-lengths of 66.0 cm and opposite currents should be placed to a distance of about 62 cm; the design was realized by connecting the coils in series and winding them in opposite directions. Only a single turn was used. According to our computations, this design should produce a field linearity of 0.8% in the volume of interest.

The design of the transverse gradient coil was based on a planar projection of a cylindrical Golay coil. The ‘Golay-type’ coils were assumed to lie on the faces of the cubical coil frame, which determined the distance of adjacent coils to 78.5 cm. The dimensions of the coils were then selected to provide sufficient field linearity while not restricting the field of view from inside the system. The selected coil dimensions and current directions are presented in Fig. 3. The individual coils comprising the gradient coil are connected in series and the number of turns is six. The coil should provide a gradient linearity of 3.0% in the volume.

After the construction of the coils, the coil fields were measured with a three-axis fluxgate magnetometer (Mag-03MC1000) by feeding sinusoidal currents into the coils and detecting the field using the lock-in technique. The fields were measured in a uniform 5 × 5 × 5 grid with overall dimensions of 17.2 × 17.2 × 15.2 cm^3^ at the center of the system. Fig. 3 represents the measured field components for the three different coil types. Within that volume, the homogeneities of the three coils were 4.0%, 3.6% and 3.6% while the field linearities of the longitudinal gradient coils were 3.0% and 3.5% and those of the transverse gradient coils were 3.5%, 3.7% and 3.6%. The homogeneous components of the homogeneous-field coils were orthogonal within 1°.

Each coil is connected to the system electronics via a 400-ohm resistor, which determines the impedance of the coil within the bandwidth of the OPM. Except of the homogeneous *B*_z_-coil, the field amplitudes of the coils are limited by the maximum output current of the electronics (around 20 mA); field amplitude of the homogeneous *B*_z_-coil is limited by the resistor. The maximum amplitudes of the homogeneous *B*_x_- and *B*_y_-coils were measured to be approximately 160 nT and that of the *B*_z_-coil about 220 nT. The maximum longitudinal and transverse gradients provided by the gradient coils were approximately 100 and 120 nT/m, respectively.

#### 2.2.2. Electronics and software

The system electronics digitize and collect the analog output of the OPM sensor electronics, transform the OPM output to control parameters that are low-pass filtered and fed to the proportional integral (PI) controllers, whose outputs are transformed and converted to analog coil voltages (linearly proportional to currents as the coil impedance is constant within the OPM bandwidth). For flexibility, the electronics include adjustable transformations (implemented as matrix multiplications) from OPM outputs to control parameters and from control parameters to the coil currents. The low-pass filter prior to the PI controller enables selecting the range of compensated frequencies (for example, to compensate only low-frequency field drifts). Schematic diagram of the electronics is presented in Fig. 4.

**Figure 4:**
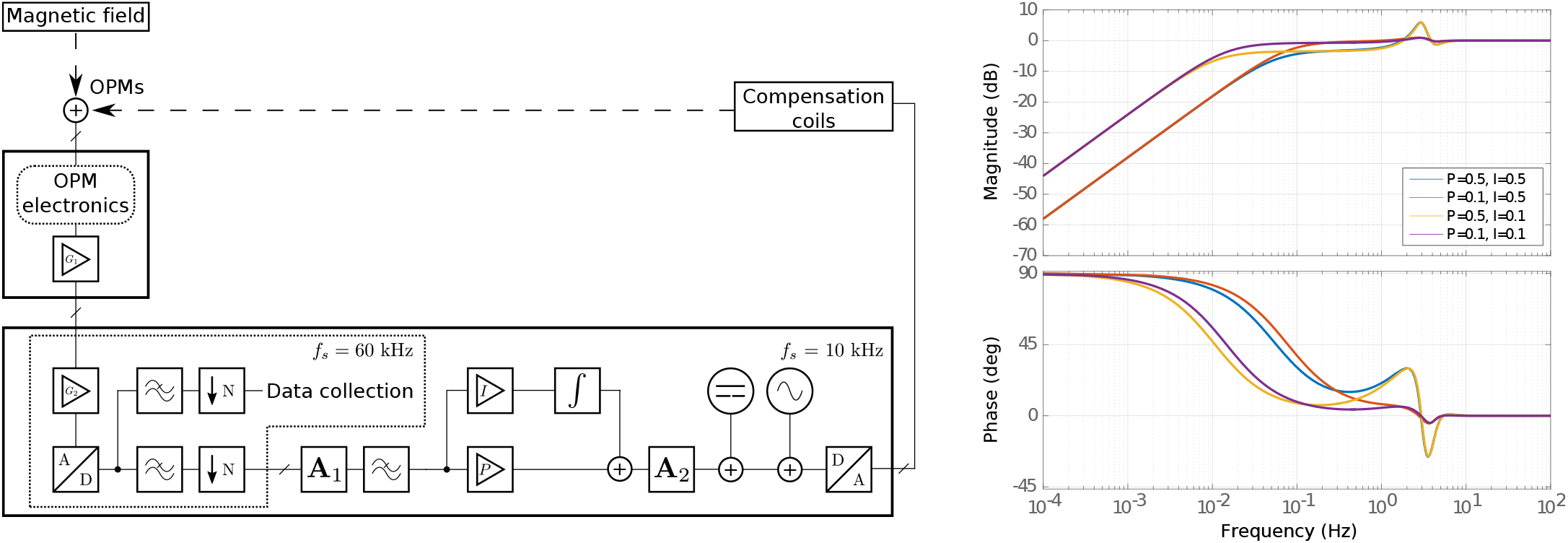
**Left:** Schematic diagram of the system electronics. OPMs measure magnetic field that comprises remanent DC field and its drifts, external interference, and the neuromagnetic field of interest. The analog output of OPM electronics is amplified, digitized and low-pass filtered to feed PI controllers whose outputs are converted to analog coil currents that zero the desired field components in the OPM array. **Right:** Bode plots of the steady-state frequency responses of the feedback loop with various *P* and *I* gains. The cut-off frequency of the low-pass filter is at 4 Hz.

We collect and digitize the analog output of the OPM sensor electronics (digital-to-analog converted at 16-bit resolution) with 12-bit analog-to-digital converters (ADCs) with a sampling frequency of 60 kHz. Two variable gains between the OPM output and the input of our electronics allow matching the voltage ranges. The gain of the OPM output can be set to 0.33, 1.0 or 3.0, and the gain of the amplifier of our electronics to 1.0 or 9.82. The digitized OPM data are then divided to two streams: one is low-pass filtered at 3.3 kHz, downsampled to a 10-kHz sampling rate, and fed to the PI controller, while the other is low-pass filtered (typ. at 330 Hz) and downsampled to the desired sampling frequency of the measurement (typ. 1 kHz) and fed to the data collection system. The initial oversampling at 60 kHz and the following low-pass filtering increase the effective resolution of the two digital signals by one or two bits.

In the negative-feedback system, the OPM signals are first multiplied by a user-defined matrix **A**_1_ that maps them to the control variables of the PI controller. Thus, different weighted combinations of the OPM signals can be used to estimate the homogeneous components and gradients of the field. Prior to the PI controller, the control variables are filtered with a low-pass filter whose cutoff frequency can be adjusted. The outputs of the PI controllers are then mapped to coil currents with a matrix multiplication by **A**_2_. Offsets are added to the coil currents for the DC compensation of the field. Additionally, sinusoidal, triangular or white noise currents can be added. Finally, the coil currents are fed to the coils through a 24-bit digital-to-analog conversion. The parameters of the electronics are controlled with a Python-based interface.

The low-pass filter in the PI loop is a sixth-order Butterworth filter realized using a delta operator in a direct-form II transposed structure to tackle finite-word-length effects in the arithmetics of the DSP (Kauraniemi et al., 1998); this filter comprises three such second-order sections. The Butterworth filter provides flat passband frequency response, relatively linear phase response in the passband, and a roll-off of 120 dB per decade. We have verified that cut-off frequencies down to 1 Hz can be realized by simulating the filter in MATLAB with fixed-point arithmetic mimicking the DSP (Iivanainen, 2016).

We simulated the steady-state transfer function of the digital feedback loop of the system using the Linear Analysis tool of the Simulink toolbox in MATLAB. Computed Bode plots of the feedback loop with the cut-off frequency of the low-pass filter at 4 Hz with multiple *P* and *I* gains of the PI controller are shown in Fig. 4. The frequency and phase responses show an anomaly near the cut-off frequency of the filter. The magnitude of this distortion increases as *P* gain increases; with a moderate *P* gain, it is less than 1 dB. The low-frequency attenuation (≤ 0.1 Hz) improves as I increases. Even if the cut-off is at 4 Hz, the attenuation approaches zero decibels well below 1 Hz with proper loop tuning (*P* = 0.1; *I* = 0.5).

#### 2.2.3. Shielding strategy

As mentioned before, we employ the same sensors that measure the neuromagnetic field for both static and dynamic shielding against the ambient field. As the sensors can be in different slots of the helmet and the position of the helmet with respect to the coils can vary between measurements, the geometric arrangement of the sensors and coils cannot be assumed fixed as in reference-array solutions. Instead, the transformation of the sensor outputs to the homogeneous and gradient fields produced by the array has to be established for every measurement. The sensor configuration in each measurement should also be such that accurate estimates of the homogeneous and gradient field can be computed.

We zero the magnetic field in the sensor array the following way. We measure the coupling matrix that describes the linear coupling of the coil currents to the magnetic field measured by the sensors by exciting the field coils and measuring the response of the sensor array to the excitation. By inverting the coupling matrix, estimates for the coil currents that zero the field in the array can be computed. For static compensation, we use the field values of the two orthogonal components that the OPM provides (dual-axis mode) while for dynamic compensation we use the OPM in a single-axis mode. After the global DC field is reduced in the sensor array with our coils, we zero the local residual field inside the sensor with the internal coils of the sensor.

## 3. Results

In this section, we present the results of the OPM measurements with the shielding system in the two-layer MSR. First, we characterize the calibration drift of the OPMs (i.e. gain and sensing-angle drifts) in the room. Second, we present the static and dynamic shielding performance of the system. Third, we quantify the OPM calibration when dynamic shielding is employed. Last, we present on-scalp somatosensory neuromagnetic responses to electric stimulation of the median nerve measured with the actively-shielded OPM array.

### 3.1. Quantification of the OPM calibration

We employed the following setup to quantify the OPM calibration drift due to the ambient field drift inside the two-layer room. We placed three OPMs in the center of the coil system so that their sensitive axes corresponded roughly to the orthogonal axes of the homogeneous coils of the system (OPMx, OPMy and OPMz). The laser beams of the OPMs were along the following axes: OPMx: z, OPMy: x and OPMz: y. The distance between any pair of sensors was at least 6 cm. Sinusoidal homogeneous fields were produced by the coils at frequencies of 55, 60, and 65 Hz. The amplitudes of the fields were set so that they were roughly the same (~12 pT). We nulled the ambient field with our shielding system; the residual local field (below 2 nT) in the sensor positions was then further nulled with the sensor-wise coils and the sensors were calibrated. After the calibration, the ambient field drifts and the coil fields were recorded for 33 minutes. The time-varying amplitudes of the homogeneous fields were estimated from the OPM signals with the lock-in technique by windowing the data to 1-s segments. Prior to the lock-in detection of the amplitude of each homogeneous field, the contributions of the other homogeneous coil fields as well as the 50-Hz power-line noise were notch-filtered (filter width 1 Hz) and the signal was band-pass filtered (width 6 Hz) around the field frequency.

The drifts of the three homogeneous components of the ambient field and the measured amplitudes of the coil fields are presented in Fig. 5. The gain in the sensitive direction of the OPM is primarily changed by the drift of the ambient field component in that direction. The drift of the ambient field component transverse to the sensitive direction of the OPM modulate the gain of the other orthogonal transverse direction; this is particularly evident in the amplitudes measured by OPMx: *B*_y_ amplitude measured by OPMx is correlated to the drift measured by OPMz and *B*_z_ amplitude is correlated to drift along y (OPMy). Altogether, the drifts in the three components of the ambient field change the gain of the OPM to the three components of the magnetic field, i.e., the ambient field drifts tilt the sensitive axis of the OPM. Fig. 5 also shows the changes in the angles of the sensitive directions of the OPMs estimated from the measured coil fields. In this 33-minute measurement, the tilts in the sensitive axes of the sensors ranged from 2° to 5°. The drift of the sensitive axis is mostly correlated with the field component orthogonal to the plane spanned by the directions of the laser beam and the sensitive axis (OPMx: y; OPMy: z; OPMz: x).

**Figure 5:**
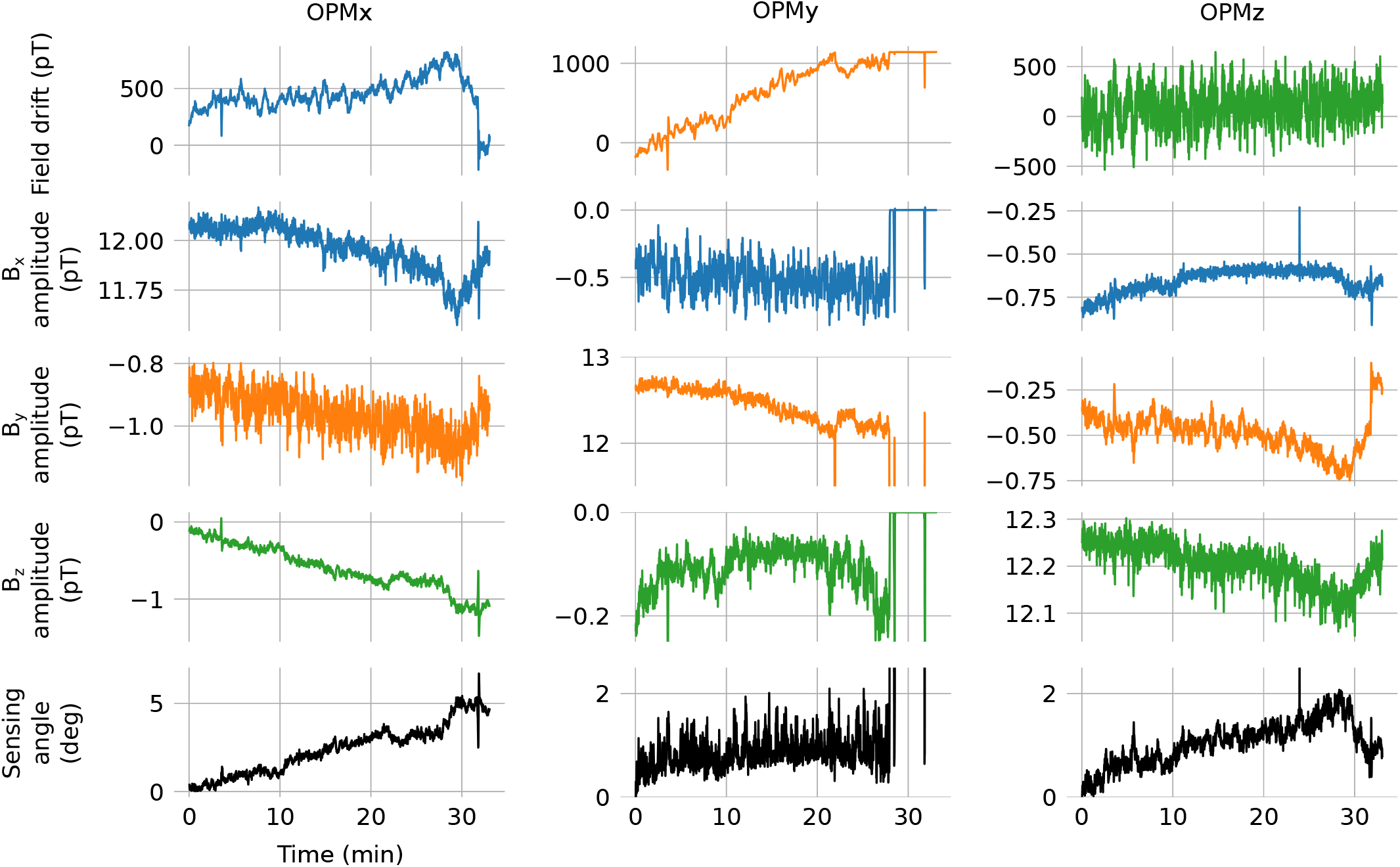
OPM calibration error due to the drift of the ambient field inside a two-layer shielded room. Three OPMs are oriented along the axes of the three homogeneous coils of the system (OPMx, OPMy and OPMz) that produce approximately orthogonal sinusoidal magnetic fields with different frequencies and roughly the same amplitudes. The measured fields in x-, y- and z-directions are colored as blue, orange and green, respectively. **First row:** The field drifts low-pass filtered below 10 Hz in the three orthogonal directions as measured by the OPMs. Near the end of the 33-min measurement the output of OPMy clips. **Rows 2–4:** The amplitudes of the three orthogonal homogeneous fields (*B*_x_, *B*_y_ and *B*_z_) as estimated with a lock-in technique. **Bottom row:** The drift of the sensitive axis of the OPM quantified as the angle that the axis makes with the initial sensing axis.

### 3.2. Performance of the shielding system

Fig. 6 shows the noise spectra of a single OPM measured in the two-layer room inside the coil system and in a three-layer shielded room (Imedco AG, Hagendorf, Switzerland) without additional shielding. In the two-layer room OPM was placed along the z-direction of the room and the coil set, which is the component with the most low-frequency drifts (see Sec. 2.1) and the highest noise from the coil set as the *B*_z_ coil has the highest gain. Spectra were estimated from five-minute recordings sampled at 1 kHz and low-pass filtered at 330 Hz using Welch’s method with half-overlapping Hanning windows of length 20 s. In the two-layer room there is considerable amount of low-frequency field noise due to the field drifts. The spectrum recorded in the two-layer room is also higher throughout the bandwidth of the OPM due to ambient magnetic noise inside the room and the noise produced by the coil system (1/*f* and white noise). At frequencies 60–80 Hz, the mean noise levels are 10.6 and 15.6 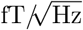 in the three- and two-layer rooms, respectively; at that band, the noise produced by the coil system is thus about 11.3 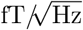.

**Figure 6:**
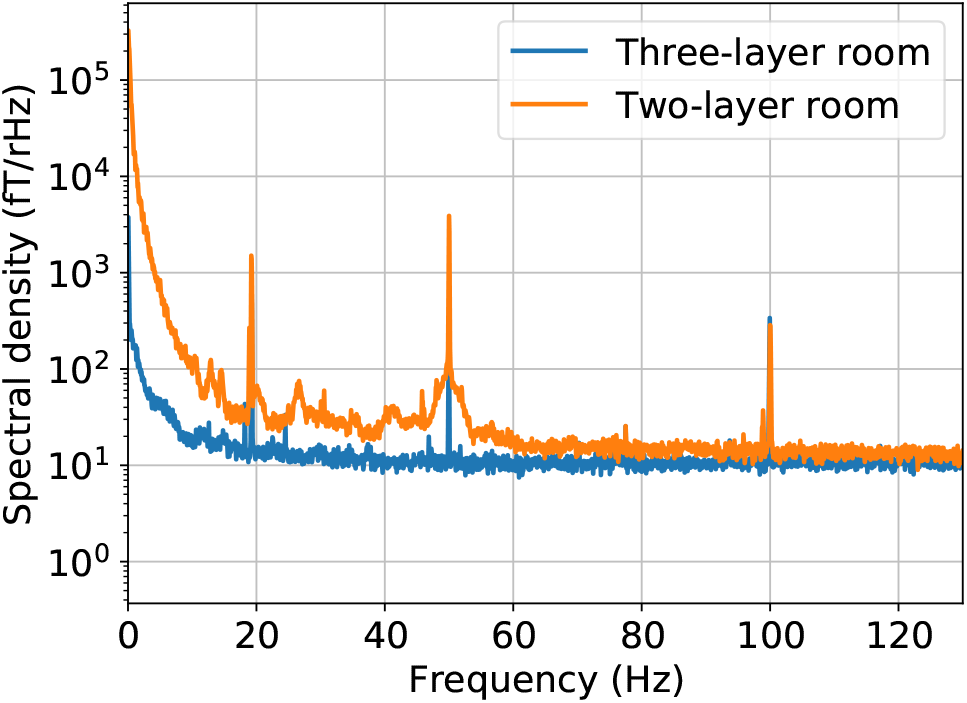
Noise spectral densities of a single OPM estimated from measurements inside a two-layer shielded room with static field compensation by the shielding system and inside a three-layer shielded room without additional shielding.

An example of the static magnetic shielding performance of our system in the two-layer room is shown in Fig. 7. The sensors were placed in a configuration that was also used to measure the somatosensory responses: six sensors are in helmet slots that cover the somatomotor cortex while two sensors are placed so that the array can measure three orthogonal components of the background field. The field was nulled using the homogeneous, longitudinal and two of the transverse gradient coils. The remanent-field components ranged from about −70 to 50 nT in the array and after nulling the components were reduced below 1 nT. The passive shielding factor of the coil system was thus about 40 dB. The maximum field drift in the room can be approximately 1 nT in ten seconds; the efficiency of the static shielding is then mostly limited by the drifts.

**Figure 7:**
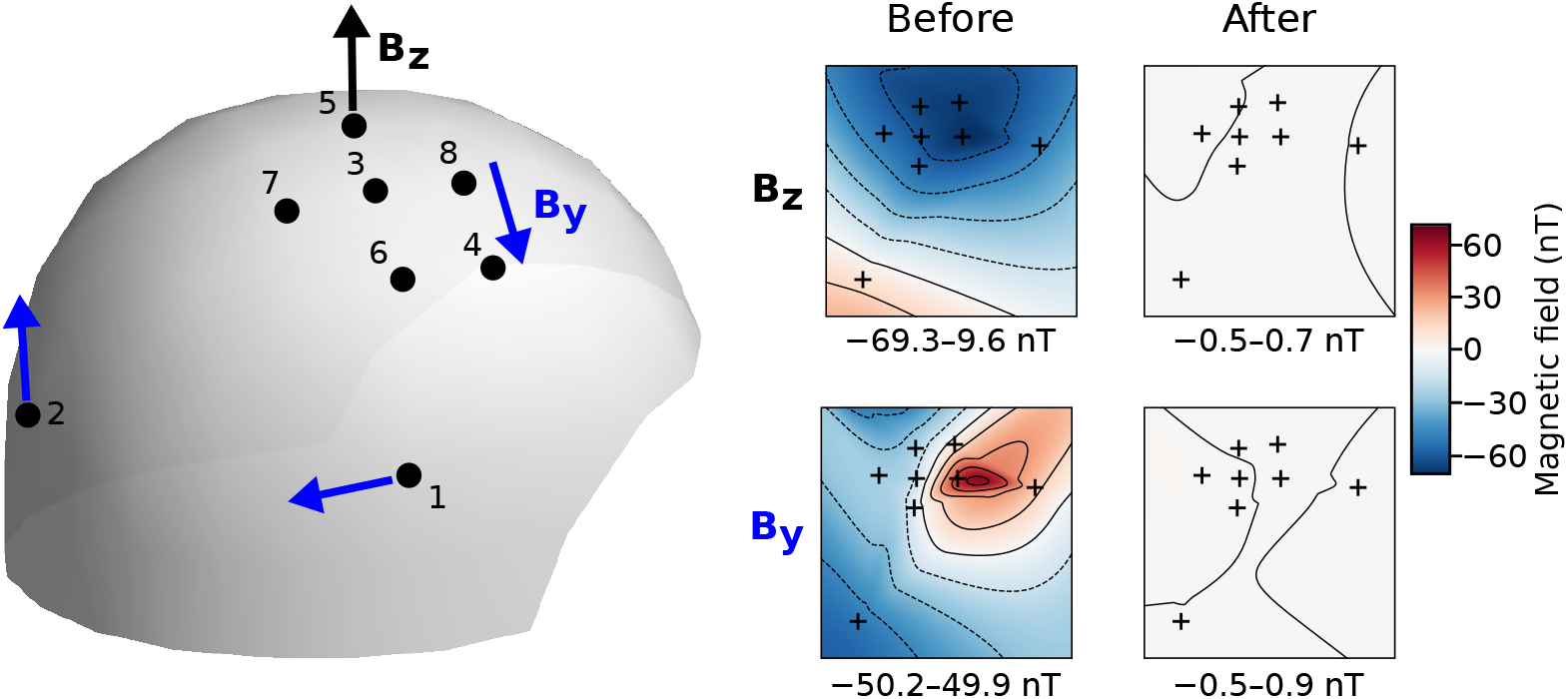
Static shielding performance of the system. **Left:** Illustration of the locations of the OPMs and the directions of their two sensitive axes. *B*_z_ denotes the out-of-the-helmet component while *B*_y_ denotes the component along the helmet surface. **Right:** The values of the field components before and after compensation with our system; the field values are visualized on a 2D-projected layout of the sensors. The ranges for the field values are shown under the field patterns.

We estimated the efficiency of the dynamic shielding of our system by comparing 15-min OPM measurements without and with dynamic shielding in the two-layer room. The sensor configuration in the measurements was the same as in Fig. 7. Before each measurement the coil system was used to reduce the static ambient field below 1.2 nT in the sensor array and the residual fields were nulled with the internal coils of the sensors. We estimated the efficiency of the dynamic shielding with two feedback loop configurations: one with low-pass filter cutoff at 4 Hz and another with no low-pass filter to measure the maximum shielding factor that can be obtained with the system. In both cases, we ran the OPM array in a negative feedback loop with the homogeneous field coils. The PI controllers were tuned by first increasing the *P* gain until the control loop started to oscillate. Then this gain was set below this critical value, and the *I* gain was increased until the loop became unstable; the *I* gain was then lowered. After the critical values were found, the *P* and *I* gains were varied below these values and the best combination of the gains was determined by visually inspecting the OPM signals, by examining the spectra of the OPM and coil outputs, and by calculating the shielding factor. Specific care was taken to avoid any resonances and to maximize the low-frequency shielding factor.

Figure 8 shows the OPM outputs (low-pass filtered below 10 Hz) with and without the dynamic compensation. Compensation removed considerable amount of low-frequency drifts of the homogeneous field components. The maximum peak-to-peak drift across the sensors during the 15-min measurement was 1.3 nT without compensation, 400 pT (filter cutoff at 4 Hz) and 30 pT (no filter) with compensation. The corresponding standard deviations were 210, 47 and 3.7 pT. The DC field offsets of the OPM outputs ranged from −90 to 40 pT with dynamic shielding. Figure 8 also shows the amplitudes of the homogeneous feedback fields and their spectra with filter cutoff at 4Hz. The field in *z*-direction had relatively larger rate of change than the *x*- and *y*-fields, as expected from the characteristics of the room (see Sec. 2.1). The shielding factors for the eight OPMs as computed from the spectra (Welch’s method; half-overlapping Hanning windows of 20 s) with and without the compensation are also shown in Figure 8. With low-pass filter cutoff at 4 Hz, the low-frequency shielding factor was on average 22 dB (range: 20.8–23.9 dB). Shielding factor reached 0 dB below 1 Hz, above which at around 4 Hz there was some distortion ranging from about −3 to 1 dB. Without the low-pass filter, the shielding factor was on average 43 dB (39.1–44.2 dB) at low frequencies and reached 0 dB approximately at 10 Hz.

**Figure 8:**
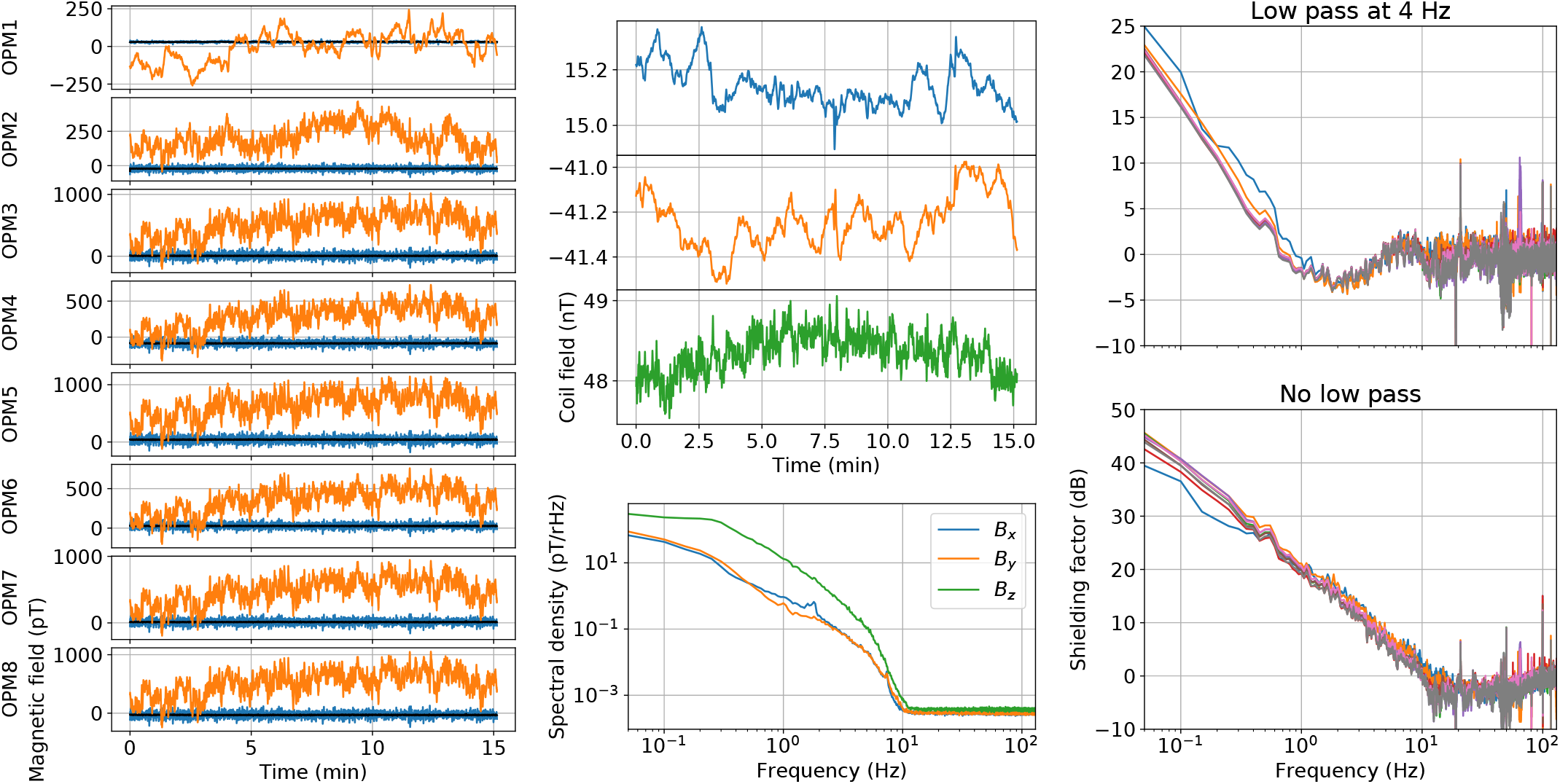
Dynamic shielding performance of the system with cutoff frequency of the negative feedback loop at 4 Hz and with no low-pass filter. **Left:** Outputs of eight OPM sensors low-pass filtered at 10 Hz without (orange) and with dynamic compensation (blue: cutoff at 4 Hz; black: no low-pass filter) during 15-min measurements. For the illustration of the sensor slots in the helmet, see Fig. 7. **Center:** The amplitudes of the feedback fields generated by the three homogeneous coils (top) and their spectra (bottom; cutoff at 4 Hz). **Right:** Shielding factor provided by the dynamic compensation as a function of frequency for the eight OPMs.

### 3.3. OPM calibration with dynamic shielding

We quantified the accuracy and drift of the OPM calibration with and without dynamic shielding by measuring the amplitude of a reference field as a function of time. The sensor configuration and PI loop tuning was the same as in Sec. 3.2. The cutoff frequency of the low-pass filter was set to 4 Hz. A 65-Hz sinusoidal field was generated with a small circular coil driven by a function generator (33120A, Hewlett Packard, Palo Alto, CA, USA). The measurements were done using two setups to estimate the calibration drift to two different reference-field shapes: in the first, the circular coil was inside the sensor helmet while in the second it was outside the helmet. The data were low-pass filtered at 330 Hz and sampled at 1 kHz. The amplitude of the reference field was computed by filtering OPM data to a 62–68-Hz band (notch at 50 Hz) and using the lock-in technique with 1-s windows. The windows where OPM signals clipped (exceeding 1.1 nT) were dropped from the analysis. In both setups, the shielded and unshielded data were cut to the same length.

The measured amplitudes of the reference signal are shown in Fig. 9 with and without dynamic shielding together with box plots summarizing the amplitude distributions. In the first measurement, there was a transient interference after five minutes lasting about ten seconds that caused the outputs of six OPMs to clip. When the reference coil was inside the helmet, the averages and standard deviations of the calibration errors (normalized by the amplitude estimate obtained by averaging across the windows) ranged across the sensors 0.1%–0.2% and 0.1%–0.15% with dynamic shielding, and 0.3%–7.6% and 0.2%–2.4% without shielding, respectively. When the coil was outside the helmet, the corresponding ranges were 0.2%–2.5% and 0.2%–1.2% with dynamic shielding while without shielding they were 1.2%–9.2% and 0.9%–7.0%, respectively. The relative error between the shielded and unshielded estimates of the reference-field amplitudes ranged across the sensors 0.3%–7.4% (coil inside) and 1.0%–23% (coil outside).

**Figure 9:**
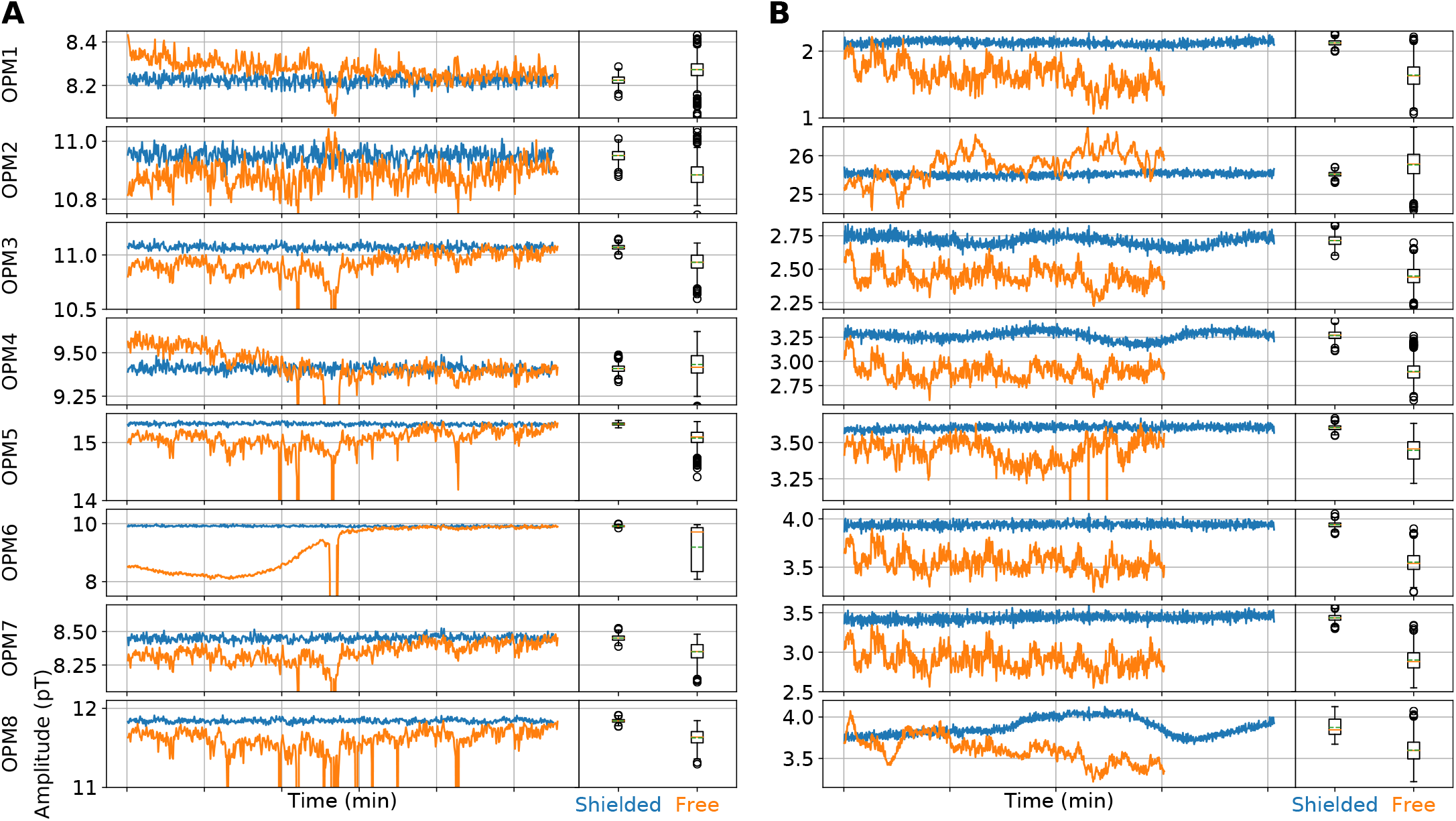
OPM calibration as quantified by the measured amplitude of a sinusoidal reference field with (blue) and without (orange) active shielding. The box plots summarize the distribution of the amplitudes. The reference field was generated with a small circular coil that was placed inside the helmet (**A**) or outside the helmet (**B**). Each time point corresponds to an estimate of the reference-signal amplitude as given by lock-in detection with a window length of 1 s. The windows in which the OPM output was clipped are discarded from the box plots. To equalize the number of samples in the box plots in B, only the amplitudes from the first 15 minutes of shielded data are included. In the box plots, the solid orange and dashed green horizontal lines indicate the median and average, respectively; the boxes extend from lower to upper quartile and the whiskers give the 1.5 interquartile ranges from the lower and upper quartiles. For the sensor locations, see Fig. 7. Data in B are the same that were used to generate Fig. 8.

### 3.4. Neuromagnetic responses to median nerve stimulation

To demonstrate the operation of the system in brain measurements, we recorded neuromagnetic responses of one subject to electric stimulation of the left median nerve at the wrist with eight dynamically-shielded OPMs. Informed consent was obtained from the subject, in agreement with the approval of the local ethics committee. The measurement setup is shown in Fig. 10A. The sensors were inserted into the same slots of the helmet as in Fig. 7 so that they touched the scalp of the subject. The sensor positions with respect to the subject’s head are shown in Fig. 10B. Before the measurement, the ambient DC field was reduced to below 1.6 nT in the OPM array and the residual local DC field was nulled with the OPMs. The OPM array was run in a negative feedback loop with the homogeneous coils using OPM outputs below 4 Hz and the same PI loop tuning as in Sec. 3.2. Prior to the measurement, about one minute of data were recorded without the subject. These data were filtered to 1–70 Hz (notch at 50 Hz), and two signal-space components (Uusitalo and Ilmoniemi (1997); see Fig. 10C) were computed to describe the signal subspace of the interference due to the compensation coils and environment. The first component is approximately homogeneous within the sensor array and most likely represents the interference from the homogeneous *B*_z_ compensation coil. The second component shows a gradient pattern.

**Figure 10:**
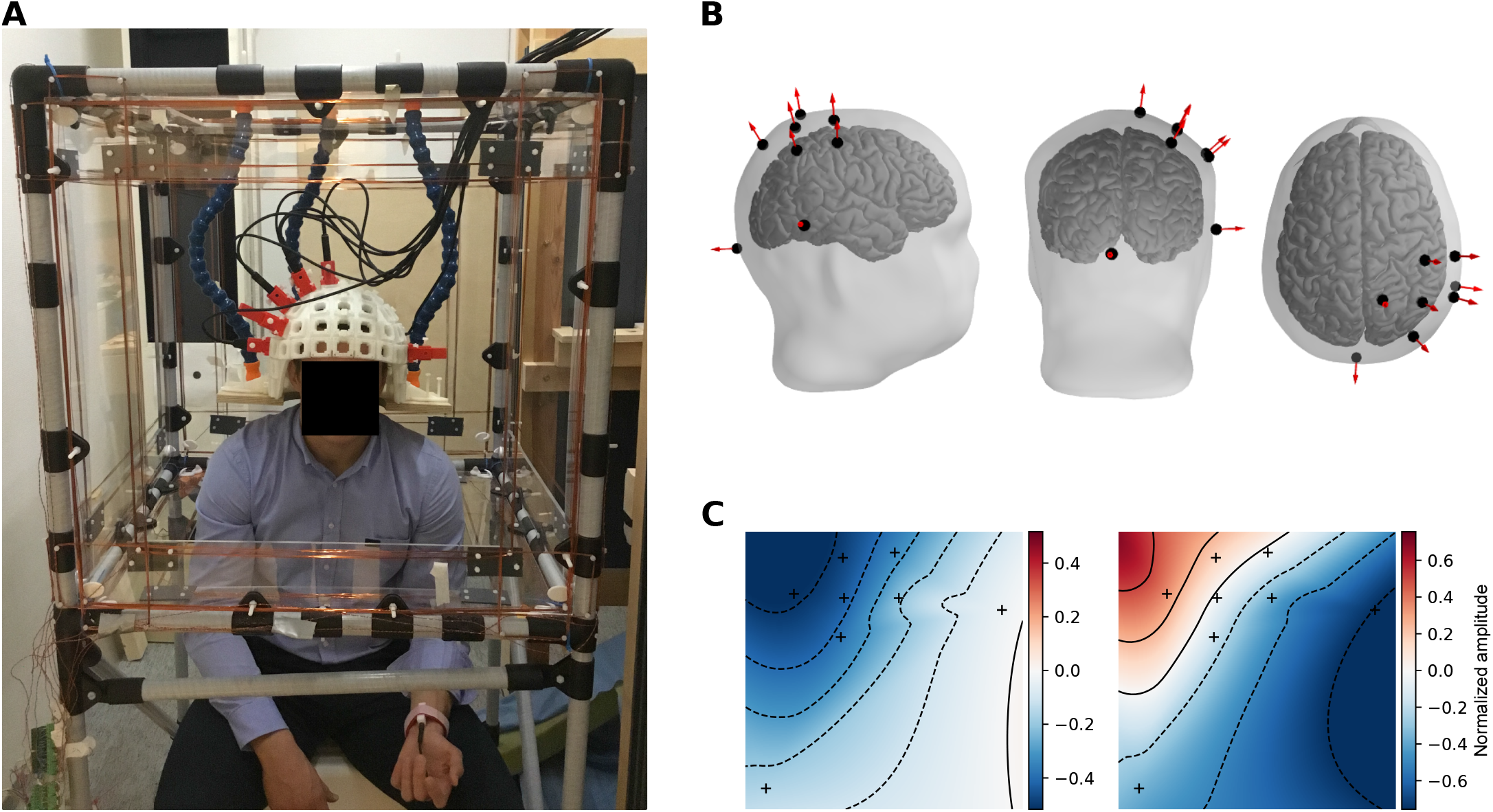
**A:** A subject inside the shielding system during the somatosensory measurement. **B:** Sensor positions with respect to the subject’s head. Six sensors covered the somatomotor cortical areas of the subject. **C:** Signal-space components describing interference subspace. The components were estimated from data measured in the absence of a subject and are visualized on a 2D-projected layout of the eight OPMs. Sensor positions are shown as black crosses.

The left median nerve of the subject was stimulated at the wrist by applying a current pulse (amplitude 7.2 mA, duration 200 *μ*s) with a current stimulator (DS7A, Digitimer Ltd, Welwyn Garden City, UK). Total number of stimuli was 153, and the inter-stimulus interval was 2.5 s with uniformly-distributed jitter of ±0.2 s. The data were band-pass filtered to 1–70 Hz (notch at 50 Hz). The two signal-space components were projected out of the data, and the mean of the baseline (0.3 s before the stimulation) amplitude was subtracted from each response. Amplitude-thresholded rejection of epochs was used: an epoch was dropped if the peak-to-peak amplitude of a single channel exceeded 7 pT during that epoch, resulting in a rejection of 10 epochs. The stimulus-locked data across the epochs were then averaged. In addition to the evoked responses, induced responses were estimated using Morlet wavelets in a frequency band of 1–49 Hz (frequency resolution: 1 Hz; number of cycles for each frequency: frequency divided by two). All analysis steps were done with the MNE software (Gramfort et al., 2014).

The evoked and induced responses are presented in Figs. 11A and 11B, respectively. Evoked responses as well as the beta-rhythm suppression and rebound around 20 Hz in the time–frequency responses are clearly visible. The evoked responses show the typical responses at latencies 22 (N20m), 41 (P40m) and 52 ms (P60m). Fig. 11D shows the dipolar field pattern of the average evoked response at three time instants corresponding to peaks at 22, 41 and 52 ms. The responses can also be seen in single trials (Fig. 11E) after signal-space projection. The amplitude of the evoked P60m-response ranged from about −1 to 1.6 pT.

**Figure 11:**
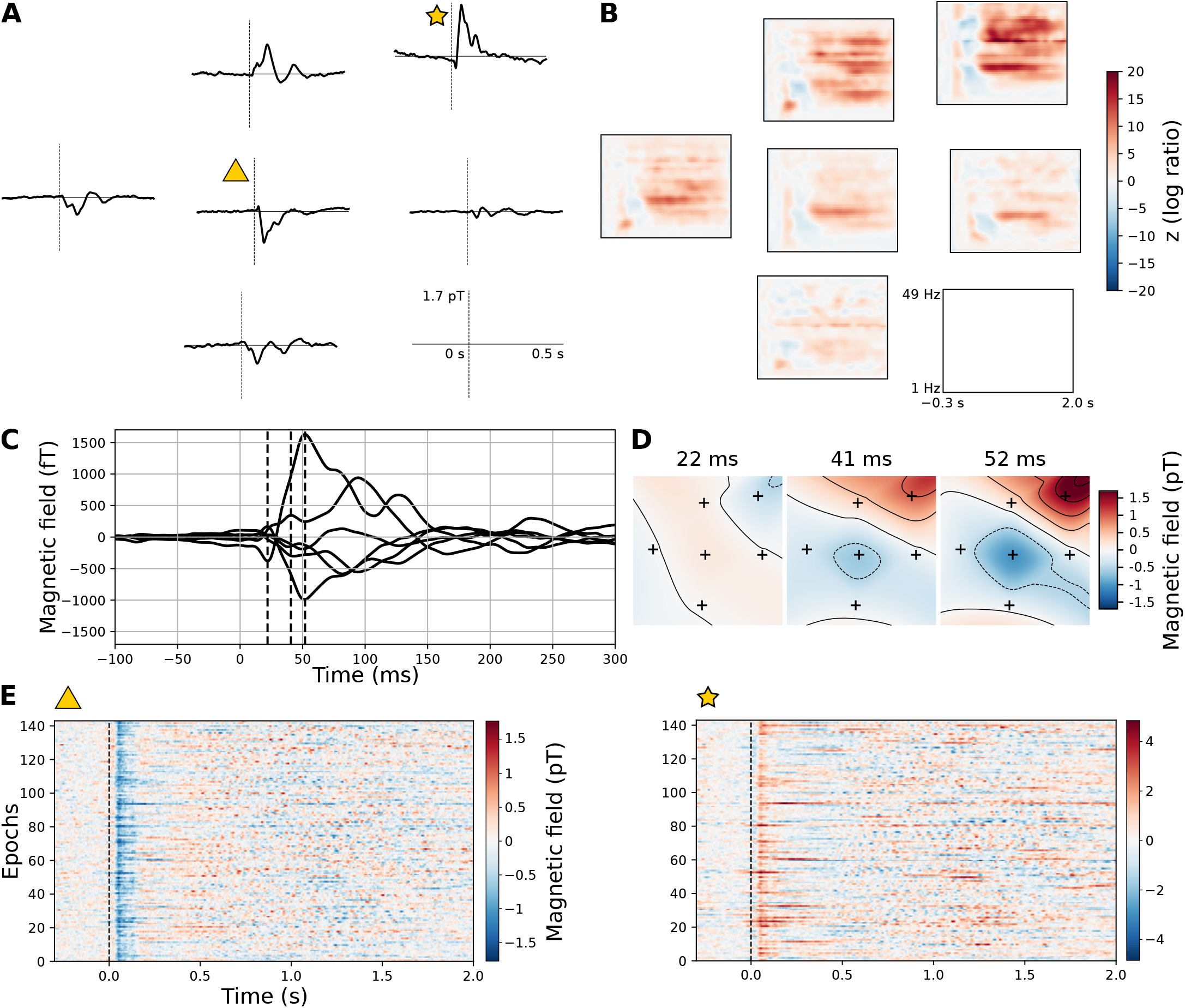
Evoked (**A**) and induced (**B**) responses to median-nerve stimulation measured with the active compensation system engaged. The responses are shown on a 2D layout of the six OPMs covering somatomotor cortex (see Fig. 10). The responses are averages of 143 trials. **C:** Butterfly plot of the average evoked responses from −100 to 300 ms. **D:** The field patterns of the response at the three time instants marked in panel C with dashed vertical lines. **E:** Single-trial responses of the sensors marked in panel A.

## 4. Discussion

We described an active field compensation system for optically-pumped magnetometers and demonstrated its use by measuring neuromagnetic fields on scalp. The system enabled the use of commercial OPMs in a two-layer MSR by bringing the remanent DC field close to zero. We emphasize that without the external DC-field compensation, these OPMs could not operate in this MSR. The dynamic shielding provided by the system considerably reduced the sensor calibration errors caused by the field drifts in the MSR. We showed that evoked and induced responses to median nerve stimulation could be detected (even without trial averaging).

Here, we discuss the obtained results and our approach to bring the OPM technology to a magnetically more challenging environment which has relatively large DC field and field drifts. Based on our findings, we try to emphasize specific points that are relevant both for the design of sensors and external field-compensation systems, to advance the development of practical OPM-based MEG systems.

### 4.1. Sensor calibration

Accurate calibration of MEG sensors is vital. Calibration error (errors in gain and in the direction of the sensitive axis) and calibration drift deteriorate the performance of source-estimation (Zetter et al., 2017), interference-removal (Taulu and Simola, 2006; Nurminen et al., 2008) and coregistration methods (Ahlfors and Ilmoniemi, 1989). For example, in the signal-space separation method for removing external interference, the knowledge of sensor calibration and cross-talk with a relative accuracy better than 1% (preferably 1 ppm) is necessary (Taulu et al., 2004, 2005).

In the OPMs that employ field modulation, changes in any of the three magnetic field components inside the sensitive volume affect the calibration of the sensor by modulating its gain and by tilting its sensitive axis (Appendix A).

In the two-layer room, the field drifts can tilt the sensitive axes of the OPMs up to about 5° (in addition to which also the gain of the OPM along that direction is changed). We have recently performed simulations to estimate the effect of the sensor position and orientations errors on the source-estimation performance of on-scalp OPM arrays (Zetter et al., 2017). We estimated that to obtain higher accuracy in source estimation with OPMs than with SQUIDs, the maximum OPM position and orientation errors should be less than 5.2 mm and 17.3°, respectively. We assumed that the errors were uniformly and independently distributed across the sensors. However, the obtained simulation results are not directly applicable here as the sensor calibration errors observed are not randomly distributed and are not independent across the sensors: the field drift causes a calibration error (gain and sensing axis error) that has a spatial dependency in the sensor array and distorts the measured field patterns and thus also the source estimates systematically. Thus, field-drift-caused calibration errors give rise to especially harmful source-estimate errors. In addition, to make most of the OPM-MEG, all the errors should be naturally minimized.

We improved the stability of OPM calibration by running the sensor array in a negative feedback loop in which a set of large coils were used to maintain a stable close-to-zero field at the sensors. This reduced the OPM calibration drift compared to unshielded operation: with and without dynamic shielding the average calibration drifts ranged from 0.1% to 2.5% and from 0.3% to 9.2%, respectively. There was also a considerable discrepancy between the unshielded reference-signal-amplitude estimate and the shielded estimate (range: 0.3%–23%) demonstrating that running the sensors without dynamic shielding leads to an error in the accuracy of the estimated field amplitude when data is averaged across time. Yet, with dynamic shielding field drifts below 4 Hz (maximum peak-to-peak drift about 400 pT), there is still some residual calibration drift originating from field drifts. In addition, in the second calibration measurement we performed there was very low-frequency and quite large calibration drift in the dynamically-shielded measurement that we believe originates from an intrinsic mechanism of the sensor.

Besides using a large external coil set to provide dynamic shielding, negative feedback to each sensor should also be employed to further improve the calibration accuracy. Internal OPM feedback has been demonstrated (Sheng et al., 2017a) but running the sensor with feedback along the sensitive axis alone does not solve the calibration issue entirely as the drifts of the transverse magnetic field components (w.r.t. sensitive axis) also affect the calibration. Therefore, an external large coil set or other means of global dynamic field control may still be needed to actively compensate also the transverse field components in the sensor positions.

In principle, deviations of sensor calibration could be measured and taken into account in addition to preventing them. The drifting calibration necessitates continuous monitoring e.g. with a high-frequency reference fields and on-or offline compensation of the calibration. However, the changing sensitive axis of the sensor complicates several analyses: e.g. in source modeling the tilting sensitive axis has to be included in the measurement model.

### 4.2. DC-field compensation

The internal coils of the OPMs should be designed and manufactured so that they can locally null magnetic field components inside typical SQUID-MEG-compatible MSRs, which can be up to ~100 nT. The 50-nT limit of the sensor used here is quite restrictive and motivated the construction of the shielding system in the first place. Even if the sensors itself could zero the local field in the MSR, the DC-field nulling with an external coil set would still be advantageous in three aspects. First, the global field nulling reduces the artefacts and calibration errors due to the sensor movement (see Sec. 2.1). Second, global nulling of the field reduces the fields that have to be generated by the internal coils of the sensors and, thereby, also the leakage of these sensor-wise DC fields to the neighboring sensors, that can broaden the resonance and shift the sensing axis of the sensor by mechanism described in Appendix A. Third, as the DC fields generated by the sensors are minimized, also the DC gradients (and other imperfections) produced to the vapor cell by the sensor-wise coils are reduced.

The homogeneous coils in our system do not have the same number of turns: there are twice as many turns in the *B*_z_-coil than in the *B*_x_- and *B*_y_-coils. The homogeneous z-component of the remanent field in the room has usually been the largest and at one point the z-component exceeded the 160-nT limit of the coils so the number of turns in the *B*_z_-coil was increased. Similar variation of the field gradients in the room has been observed. This is partly due to the changing magnetization of the MSR walls resulting from prepolarization pulses applied by the MEG-MRI device also present in the room.

The maximum field amplitude the *B*_z_-coil can produce with current electronics is about 220 nT (or 156 nT_rms_) while the noise originating from the homogeneous coils is about 11 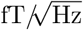 (the noise from the gradient coils is negligible). Thus, the dynamic range of the homogeneous coil is about 140 dB, which is limited by the noise from the electronics (mainly from the digital-to-analog converter). If larger ambient fields are to be compensated (e.g. in environments with very light shielding) by increasing the number of windings in the coils, the noise from the coils will begin to limit the sensitivity of the measurement. However, by reducing the voltage and current noise from the electronics the dynamic range can be increased.

### 4.3. Mixing of the neuromagnetic and shielding fields

We low-pass filter the OPM outputs to get the control variables for compensation. The low-pass filter is used to separate the brain responses from the low-frequency field drifts on individual sensor level. Here, we used a low-pass cutoff at 4 Hz, which together with the PI tuning resulted in a shielding factor that was about 22 dB at low frequencies, approached 0 dB at 1 Hz and showed some distortion around 4 Hz. The shielding fields then interfere with the neuromagnetic fields below 1 Hz and cause some distortion around 4 Hz. However, the shielding fields only cause interference with specific field topographies (in our case three orthogonal homogeneous field patterns): thus, only those components of the neuromagnetic field are affected. We attempted to use a cutoff frequency less than 4 Hz (ideally below 1 Hz) but we could not achieve satisfactory shielding with those very low cutoff frequencies. We believe that this is due to the relatively high rate-of-change of the z-component of the ambient field inside the room. Depending on the spectral characteristics of the field drifts, smaller cutoff frequencies could be used in other shielded rooms.

By high-pass filtering the obtained neuromagnetic data above 1 Hz most of the contribution of shielding fields will be removed. In addition, part of the contribution of the shielding field to the measurement was reduced by projecting out two signal-space components out of the data (the other describing the interference from the *B*_z_-coil), which increased the signal-to-noise ratio considerably. These projections also affect the measured neuromagnetic field with our 8-channel OPM array as the angles between the interference and neuromagnetic components are small in the signal space; the projection will then “eat” some of the neuromagnetic field as well.

The use of the low-pass filter is intended to be temporary. As the number of OPMs in the sensor array increases, we can use methods that take advantage of the spatial sampling provided by the array to separate the brain responses from interference (due to the coils and environment) in the signal space. With the use of such methods, the low-pass filter in the feedback loop becomes unnecessary and the uncompromised shielding performance of the system can be used reaching approximately 43 dB in the low frequencies. Additionally, the gradient coils can be also used for active shielding. Signal-space components describing the fields from all of the coils could be measured and projected out of the data; the high number of channels will then allow better separation of the shielding and neuromagnetic fields as the subspace angles increase between those. In addition, methods such as signal-space separation (Taulu and Simola, 2006) could be used to separate the neuromagnetic and interference fields. We stress that these methods necessitate accurate and steady calibration of the sensor array.

### 4.4 Wearable or rigid sensor array?

Previously, a wearable OPM sensor array has been demonstrated together with a large coil set in an MSR (homogeneous field ~20 nT; gradients ~10 nT/m; Boto et al. 2018). This system applies six bi-planar coils to null the homogeneous components and three gradients of the remanent DC field (Holmes et al., 2018). The coils produce an open and accessible environment for the subject — an attractive feature that requires a more complex coil design than simple rectangular coils. In that system, the outputs of four reference OPMs are used to null the remanent DC field before the measurement. The reference OPMs are placed at some distance from the head at fixed positions with respect to the coils. The sensor array is wearable in a sense that the subjects can move their head (and the attached sensors) during the measurements. Using the system in their shielded room, rotation of the head by ±34° and translation by ±9.7 cm led to a variation of magnetic field by 1 nT which could be further reduced to 40 pT by a linear regression of field changes correlated with head-motion parameters recorded with a camera system (Holmes et al., 2018). Such movements leading to an 1-nT artefact should also produce an OPM calibration error which the linear regression cannot remove; the calibration error will actually limit the performance of the linear regression.

Our system comprises a semi-rigid sensor array within the active shielding coil set and employs the same sensors that measure brain signals also for controlling the shielding coils. The sensor helmet is attached to the coil set with a flexible support system that allows small movements of the helmet together with the subject’s head (a snug fit of the helmet to the head is achieved by adjusting the depth of the real and additional dummy sensors). The system was built to provide static and dynamic compensation, allowing the use of the OPM sensor array in our lower-quality shielded room: DC-field compensation together with local field nulling with the sensor-wise coils bring the OPMs to their dynamic range and allows minor movements of the head and helmet while dynamic compensation reduces OPM calibration errors caused by the field drifts.

To enable robust use of wearable OPM arrays in the two-layer room is challenging because of the large homogeneous components and gradients of the field, and the large low-frequency drifts. One solution to bring a wearable array into this room would be to use a reference-sensor array to null also the temporal field drifts. The performance of such system will be determined by how well the field in the reference array estimate the field and its drifts in the measurement array. In light magnetic shields, the field can be complex and it is not guaranteed that such system will work satisfactorily. By contrast, when the shielding is driven by sensors in the measurement volume, the shielding factor should be better than in the reference-array solutions (Taulu et al., 2014). The usage of reference sensors in our two-layer room is further complicated by practical constraints: the current OPMs cannot be made sensitive without external DC-field compensation. Thus, additional coils should be used for the reference-sensor array or the present coils should be made larger and/or re-designed so that their homogeneous and linear volumes include both measurement- and reference-sensor arrays. In addition, the construction of wearable OPM arrays is also complicated by the dimensions of the OPM used here: the OPM needs comprehensive mechanical support (such as the 3D-printed headcasts; Boto et al. 2018) to stay in place on the subject’s scalp.

The concept of wearable MEG is attractive as it enables new research paradigms which include subject movement and examination of new subject groups (such as children) whose responses have been difficult to measure with SQUID-based systems. With future advances in sensor technology (e.g. reduction of sensor size) and in both passive and active shielding systems, wearable OPM systems could be potentially used even in lighter magnetic shields. Currently, we find it an intriguing task to extend our system to wearable OPM arrays that work accurately in the two-layer room.

We point out that presently the choice between rigid or wearable sensor array should always depend on the research question and the desired quality of the data. Sensor movement leads to artefacts in the data, and depending on the amplitudes of the field variations also sensor calibration can be affected. The spectral characteristics of the artefacts and whether the artefacts interfere in the frequency band of interest depend on the velocity of the head movement and how much the head ‘vibrates’. Based on our experience with wearable OPM arrays in the three-layer MSR of Aalto University (homogeneous field components ~10 nT; gradients ~6 nT/m), the movement artefacts typically contaminate low-frequency data (below ~10 Hz). If the neuromagnetic signals of interest are in the low frequencies and the task or subject population does not involve or necessitate movement, measurement with a rigid sensor array is currently the most attractive solution to ensure the best data quality.

## 5. Conclusions

We have constructed an active-shielding system to be used in on-scalp neuromagnetic measurements with OPMs and demonstrated its operation. With the system, we could measure neuromagnetic responses with OPMs in a lower-quality magnetically shielded room designed to be used with SQUID-MEG. We have taken a step forward to bring OPM-based MEG to lower-quality and even lightweight magnetically shielded rooms.

## Conflict of interest

The authors declare that they have no conflict of interest.

## Acknowledgements

The authors thank Svenja Knappe and Antti Makinen for discussions, and Koos Zevenhoven for providing the Python interface to control the data-acquisition system. Research reported in this publication received funding from Finnish Cultural Foundation (JI), the European Research Council under ERC Grant Agreement n. 678578 and the National Institute of Neurological Disorders and Stroke of the National Institutes of Health under Award Number R01NS094604. The content is solely the responsibility of the authors and does not necessarily represent the official views of the funding organisations.

## Appendix A. Calibration of an OPM

Assuming that the background magnetic field is nulled, the output of an optically-pumped magnetometer utilizing field modulation in z-direction (i.e. it is sensitive in that direction) and probing atoms with a laser beam in x-direction is proportional to the spin-polarization in x-direction (Cohen-Tannoudji et al., 1970; Slocum and Marton, 1973; Shah and Romalis, 2009)

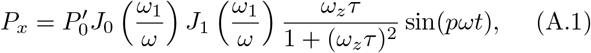

where 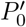 is related to the steady-state polarization 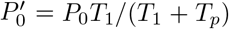, *J_n_* is a Bessel function of order *n, ω*_1_ = *γB*_1_ is the Larmor frequency of the modulation field *B*_1, *ω*_ is the frequency of the modulation field, *ω_z_* is the Larmor frequency of the field that is detected, *p* is an odd integer and *τ* is the atomic spin coherence time given by

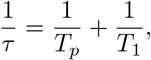

where 1/*T_p_* is the optical pumping rate and *T*_1_ is the relaxation time of the atoms. The dispersive lineshape of Eq. A.1 can be extracted by demodulating at *ω* assuming *p* = 1.

When transverse magnetic field components (*ω_x_* and *ω_y_*) are present, the output of the magnetometer is given by the series (Cohen-Tannoudji et al., 1970)

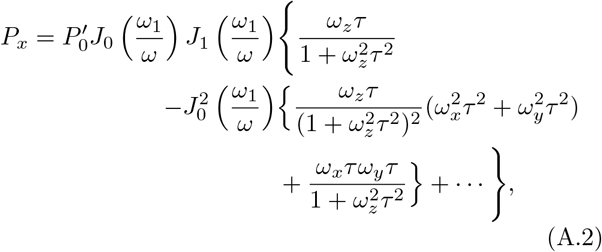

where it is assumed that *ω_x_τ, ω_y_τ* ≪ 1. Assuming that the magnetic field consists of large low-frequency drifting components (*B_x_, B_y_, B_z_*) and smaller components presenting the signal of interest (*δB_x_, δB_y_, δB_z_*), i.e., making the following substitutions to Eq. A.2

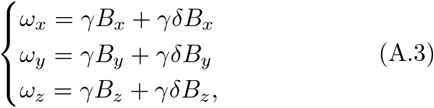

where *B_x_* ≫ *δB_x_, B_y_* ≫ *δB_y_* and *B_z_* ≫ *δB_z_*, Eq. A.2 can be simplified and approximated to (by neglecting small terms and low-frequency drift terms)

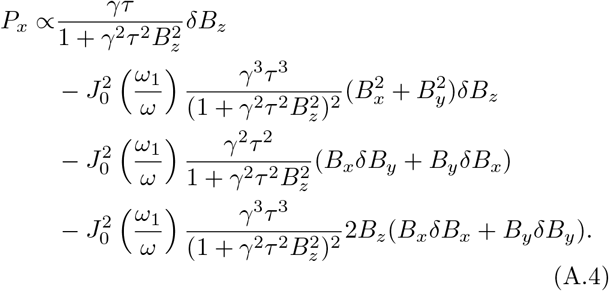

The first term in Eq. A.4 describes the effect of longitudinal field drifts to the gain of the sensor (derived also in the supplements of the study by Boto and colleagues (2018)) while the other terms describe the effects of the transverse-field-component drifts to the sensor output. The second term shows that the transverse components modulate the gain of the sensor in a second order. The third term shows that the drifts of the transverse components make the sensor sensitive also to the transverse components of the field of interest: field drift in x-direction makes the sensor sensitive to y-direction and vice versa. The last term describes the coupled second-order effect of longitudinal and transverse field drifts. Altogether, the field drifts affect the sensor calibration by changing the gain and tilting the sensitive axis of the sensor.

## References

Ahlfors, S. and Ilmoniemi, R. J. (1989). Magnetometer position indicator for multichannel MEG. In Advances in biomagnetism, pages 693–696. Springer.

Alem, O., Mhaskar, R., Jiménez-Martínez, R., Sheng, D., LeBlanc, J., Trahms, L., Sander, T., Kitching, J., and Knappe, S. (2017). Magnetic field imaging with microfabricated optically-pumped magnetometers. Optics Express, 25(7):7849–7858.

Allred, J., Lyman, R., Kornack, T., and Romalis, M. (2002). High-sensitivity atomic magnetometer unaffected by spin-exchange relaxation. Physical review letters, 89(13):130801.

Baillet, S. (2017). Magnetoencephalography for brain electrophysiology and imaging. Nature neuroscience, 20(3):327.

Borna, A., Carter, T. R., Goldberg, J. D., Colombo, A. P., Jau, Y.-Y., Berry, C., McKay, J., Stephen, J., Weisend, M., and Schwindt, P. D. (2017). A 20-channel magnetoencephalography system based on optically pumped magnetometers. Physics in Medicine & Biology, 62(23):8909.

Boto, E., Bowtell, R., Krüger, P., Fromhold, T. M., Morris, P. G., Meyer, S. S., Barnes, G. R., and Brookes, M. J. (2016). On the potential of a new generation of magnetometers for MEG: a beam-former simulation study. PloS one, 11(8):e0157655.

Boto, E., Holmes, N., Leggett, J., Roberts, G., Shah, V., Meyer, S. S., Muñoz, L. D., Mullinger, K. J., Tierney, T. M., Bestmann, S., et al. (2018). Moving magnetoencephalography towards real-world applications with a wearable system. Nature.

Budker, D. and Romalis, M. (2007). Optical magnetometry. Nature Physics, 3(4):227.

Cohen-Tannoudji, C., Dupont-Roc, J., Haroche, S., and Laloё, F. (1970). Diverses résonances de croisement de niveaux sur des atomes pompés optiquement en champ nul. I. Théorie. Revue de physique appliquée, 5(1):95–101.

Colombo, A. P., Carter, T. R., Borna, A., Jau, Y.-Y., Johnson, C. N., Dagel, A. L., and Schwindt, P. D. (2016). Four-channel optically pumped atomic magnetometer for magnetoencephalography. Optics express, 24(14):15403–15416.

Ditterich, J. and Eggert, T. (2001). Improving the homogeneity of the magnetic field in the magnetic search coil technique. IEEE transactions on biomedical engineering, 48(10):1178–1185.

Faley, M., Dammers, J., Maslennikov, Y., Schneiderman, J., Winkler, D., Koshelets, V., Shah, N., and Dunin-Borkowski, R. (2017). High-Tc SQUID biomagnetometers. Superconductor Science and Technology, 30(8):083001.

Gramfort, A., Luessi, M., Larson, E., Engemann, D. A., Strohmeier, D., Brodbeck, C., Parkkonen, L., and Hämäläinen, M. S. (2014). MNE software for processing MEG and EEG data. NeuroImage, 86:446–460.

Hämäläinen, M., Hari, R., Ilmoniemi, R. J., Knuutila, J., and Lounasmaa, O. V. (1993). Magnetoencephalography–theory, instrumentation, and applications to noninvasive studies of the working human brain. Reviews of Modern Physics, 65(2):413.

Holmes, N., Leggett, J., Boto, E., Roberts, G., Hill, R. M., Tierney, T. M., Shah, V., Barnes, G. R., Brookes, M. J., and Bowtell, R. (2018). A bi-planar coil system for nulling background magnetic fields in scalp mounted magnetoencephalography. NeuroImage.

Iivanainen, J. (2016). Active magnetic shield for optical neuromagnetic measurements. M. Sc. thesis, Aalto University.

Iivanainen, J., Stenroos, M., and Parkkonen, L. (2017). Measuring MEG closer to the brain: Performance of on-scalp sensor arrays. NeuroImage, 147:542–553.

Johnson, C. N., Schwindt, P., and Weisend, M. (2013). Multi-sensor magnetoencephalography with atomic magnetometers. Physics in Medicine & Biology, 58(17):6065.

Kauraniemi, J., Laakso, T. I., Hartimo, I., and Ovaska, S. J. (1998). Delta operator realizations of direct-form IIR filters. IEEE Transactions on Circuits and Systems II: Analog and Digital Signal Processing, 45(1):41–52.

Nurminen, J., Taulu, S., and Okada, Y. (2008). Effects of sensor calibration, balancing and parametrization on the signal space separation method. Physics in Medicine & Biology, 53(7):1975.

Osborne, J., Orton, J., Alem, O., and Shah, V. (2018). Fully integrated, standalone zero field optically pumped magnetometer for biomagnetism. In Steep Dispersion Engineering and Opto-Atomic Precision Metrology XI, volume 10548, page 105481G. International Society for Optics and Photonics.

Riaz, B., Pfeiffer, C., and Schneiderman, J. F. (2017). Evaluation of realistic layouts for next generation on-scalp MEG: spatial information density maps. Scientific Reports, 7(1):6974.

Shah, V. and Romalis, M. (2009). Spin-exchange relaxation-free magnetometry using elliptically polarized light. Physical Review A, 80(1):013416.

Sheng, D., Perry, A. R., Krzyzewski, S. P., Geller, S., Kitching, J., and Knappe, S. (2017a). A microfabricated optically-pumped magnetic gradiometer. Applied physics letters, 110(3):031106.

Sheng, J., Wan, S., Sun, Y., Dou, R., Guo, Y., Wei, K., He, K., Qin, J., and Gao, J.-H. (2017b). Magnetoencephalography with a Cs-based high-sensitivity compact atomic magnetometer. Review of Scientific Instruments, 88(9):094304.

Slocum, R. and Marton, B. (1973). Measurement of weak magnetic fields using zero-field parametric resonance in optically pumped He 4. IEEE Transactions on Magnetics, 9(3):221–226.

Taulu, S., Kajola, M., and Simola, J. (2004). Suppression of interference and artifacts by the signal space separation method. Brain topography, 16(4):269–275.

Taulu, S. and Simola, J. (2006). Spatiotemporal signal space separation method for rejecting nearby interference in MEG measurements. Physics in Medicine & Biology, 51(7):1759.

Taulu, S., Simola, J., and Kajola, M. (2005). Applications of the signal space separation method. IEEE transactions on signal processing, 53(9):3359–3372.

Taulu, S., Simola, J., Nenonen, J., and Parkkonen, L. (2014). Novel noise reduction methods. In Magnetoencephalography, pages 35–71. Springer.

Uusitalo, M. A. and Ilmoniemi, R. J. (1997). Signal-space projection method for separating MEG or EEG into components. Medical and Biological Engineering and Computing, 35(2):135–140.

Vesanen, P. T., Nieminen, J. O., Zevenhoven, K. C., Dabek, J., Parkkonen, L. T., Zhdanov, A. V., Luomahaara, J., Hassel, J., Penttilä, J., Simola, J., et al. (2013). Hybrid ultra-low-field MRI and magnetoencephalography system based on a commercial whole-head neuromagnetometer. Magnetic Resonance in Medicine, 69(6):1795–1804.

Zetter, R., Iivanainen, J., Stenroos, M., and Parkkonen, L. (2017). Requirements for coregistration accuracy in on-scalp MEG. Brain Topography, pages 1–18.

